# Joint Inference of Clonal Structure using Single-cell Genome and Transcriptome Sequencing Data

**DOI:** 10.1101/2020.02.04.934455

**Authors:** Xiangqi Bai, Zhana Duren, Lin Wan, Li C. Xia

## Abstract

Latest advancements in high-throughput single-cell genome (scDNA) and transcriptome (scRNA) sequencing technologies enabled cell-resolved investigation of tissue clones. However, it remains challenging to cluster and couple single cells for heterogeneous scRNA and scDNA data generated from the same specimen. In this study, we present a computational framework called CC-NMF, which employs a novel Coupled-Clone Non-negative Matrix Factorization technique to jointly infer clonal structure for matched scDNA and scRNA data. CCNMF couples multi-omics single cells by linking copy number and gene expression profiles through their general concordance. We validated CC-NMF using both simulated benchmarks and real-world applications, demon-strating its robustness and accuracy. We analyzed scRNA and scDNA data from an ovarian cancer cell lines mixture, a gastric cancer cell line, as well as a primary gastric cancer, successfully resolving underlying clonal structures and identifying high correlations of coexisting clones between genome and transcriptome. Overall, CCNMF is a coherent computational framework that simultaneously resolves genomic and transcriptomic clonal architecture, facilitating understanding of how cellular gene expression changes along with clonal genome alternations.

## Introduction

Understanding of how genomic content changes impact gene expression in individual cells is essential to further understand the cell clone development in normal and diseased tissues. In particular, characterizing the clone-wise gene dosage effect, i.e., the sensitivity of cellular gene expression to the copy number profile shared by the group of cells, is critical to elucidate the functional consequence of diseases-associated genomic copy number variants (CNVs), a significant challenge in current structural variant research (*1–3*).

However, there is no technology available that can efficiently and accurately measure both DNA copy number and gene expression profiles of individual cells simultaneously. Although several technologies (*4–6*), like scTrio-seq had made an attempt to measure genomic and transcriptomic content of up to a few cells per batch, it remains a low-throughput technique. Highthroughput single-cell sequencing technologies that are currently available can only measure either the transcriptome (*7–10*) or the genome (*11–13*) content of individual cells, but not both simultaneously.

For examples, droplet-based single-cell RNA sequencing (scRNA) technology is routinely employed to measure cellular expression so as to assess the clonal development states of various tissues and cell systems (*12, 14*). Recently, droplet-based single-cell DNA sequencing (scDNA) technology enabled cell-wise and genome-wide measurement of genomic alternations, such as copy number variants, in thousands of cells (*12, 15*). High quality single-cell copy number variants combined with single-cell gene expression profiles promise to further reveal the clonal heterogeneity in complex tissues and cell systems (*16–20*). Realizing the potential, however, will require high fidelity co-clustering of heterogeneous single cells measured scRNA and scDNA sequencing technologies.

Addressing this difficulty, we developed an efficient computational method – **C**oupled-**C**lone **N**on-negative **M**atrix **F**actorization, termed as CCNMF. It reasonably models the shared underlying clonal structure and the general concordance between cellular expression level and copy number states (*12, 21*). CCNMF then employs machine learning algorithms to infer the most likely multi-modal integration solution. Briefly, CCNMF took two matrices as inputs: single-cell gene expression matrix obtained through scRNA-seq technology and single-cell copy number variants matrix obtained from scDNA-seq technology, both derived from the same biological specimen.

CCNMF was established on a model-based approach which couples single cells across scDNA and scRNA data by maximizing their global concordance between gene expression and copy number variants. CCNMF optimizes an objective function that simultaneously maximizes intra-clone compactness and inter-clone structure coherence, this coupled-clone nonnegative matrix factorization framework followed the co-clustering concept as introduced in ref. (*22*). A coherent underlying clonal structure, i.e. the identity-linked cell clusters between scDNA and scRNA data, is thus inferred as the weight matrix optimally assigning all cells to their most likely cluster identity. Based on that, CCNMF accurately estimates the dosage effect per gene and cell cluster.

Before CCNMF, only a few methods were available for analyzing the combined scDNA and scRNA data. These methods mostly operate in a map-to-reference mechanism, i.e., data from one technology is mapped to the reference clonal structure derived from another technology (*21, 23–27*). For examples, *clonealign*, an early attempt of integrative modeling gene dosage effect of DNA copy number and gene expression, statistically assigns scRNA gene expression states to a reference phylogenetic tree representing scDNA-derived clones, in a Bayesian way (*21*); *Seurat*, which mainly integrates multiple scRNA datasets, can also project other types of single cell data to the scRNA-derived clusters using the mutual nearest neighbor search (*23*); *DENDRO* infers single cell copy numbers from scRNA data and validates the result using the scDNA data (*24*). The model-free single-cell integrative nonnegative matrix factorization (iNMF) method (*25*) identified the shared cell types across multi samples and species from scRNA-seq data. However, these map-to-reference inference methods risk systematic bias because the choice of reference technology was largely arbitrary, and different choices significantly influence downstream analysis.

Instead, CCNMF takes a data-driven approach, unbiased toward data and technology sources. CCNMF utilizes the coherence of the underlying clonal structure shared within the biological specimen to maximize the inference for true cell clonal identity and gene expression effects. Using both simulated and real cancer datasets, we validated that CCNMF can faithfully recover the underlying clonal structure, accurately identify clonal identity for all single cells, and statistically infer gene-wise dosage and expression changes that differentiate each clone. We applied CCNMF to characterize an ovarian cancer cell mixture, a gastric cancer cell line and a primary gastric cancer, and the results showed CCNMF is capable of identifying clonal structure and dosage effect in real cell systems. Thus, scDNA-and scRNA-seq combined with CCNMF analysis offers a new way to study the functional consequence of clonal gene dosage change and how it contributes to clonal development.

## Results

### The CCNMF Toolkit

The CCNMF analytical framework has been implemented as an *R* package, with a workflow illustrated in Fig. 1. CCNMF toolkit can accept scRNA and scDNA data in standard formats, including those generated by 10X Genomics scRNA-/scDNA-seq and DLP scDNA technologies. The toolkit executes the statistical framework and analytical steps as follows:

1. Aligns scRNA genes and scDNA genome bins to the same genome reference.
2. One-to-multiple mapping of scDNA bins to genes using location overlapping (*>* 1bp) to reduce the scDNA data to the same gene coordinates as the scRNA data.
3. Initializes the coupled term between scRNA (*E*) and scDNA data (*O*) using a coupling matrix (*A*), which is either an identity matrix or a user provided matrix with prior information.
4. Iteratively optimizes the objective function using a gradient descent algorithm until convergence.
5. Identifies the most coherent clonal structure by finding the maximum weights of the *H* matrices that represented the most likely cell clonality membership.

**Fig. 1.**
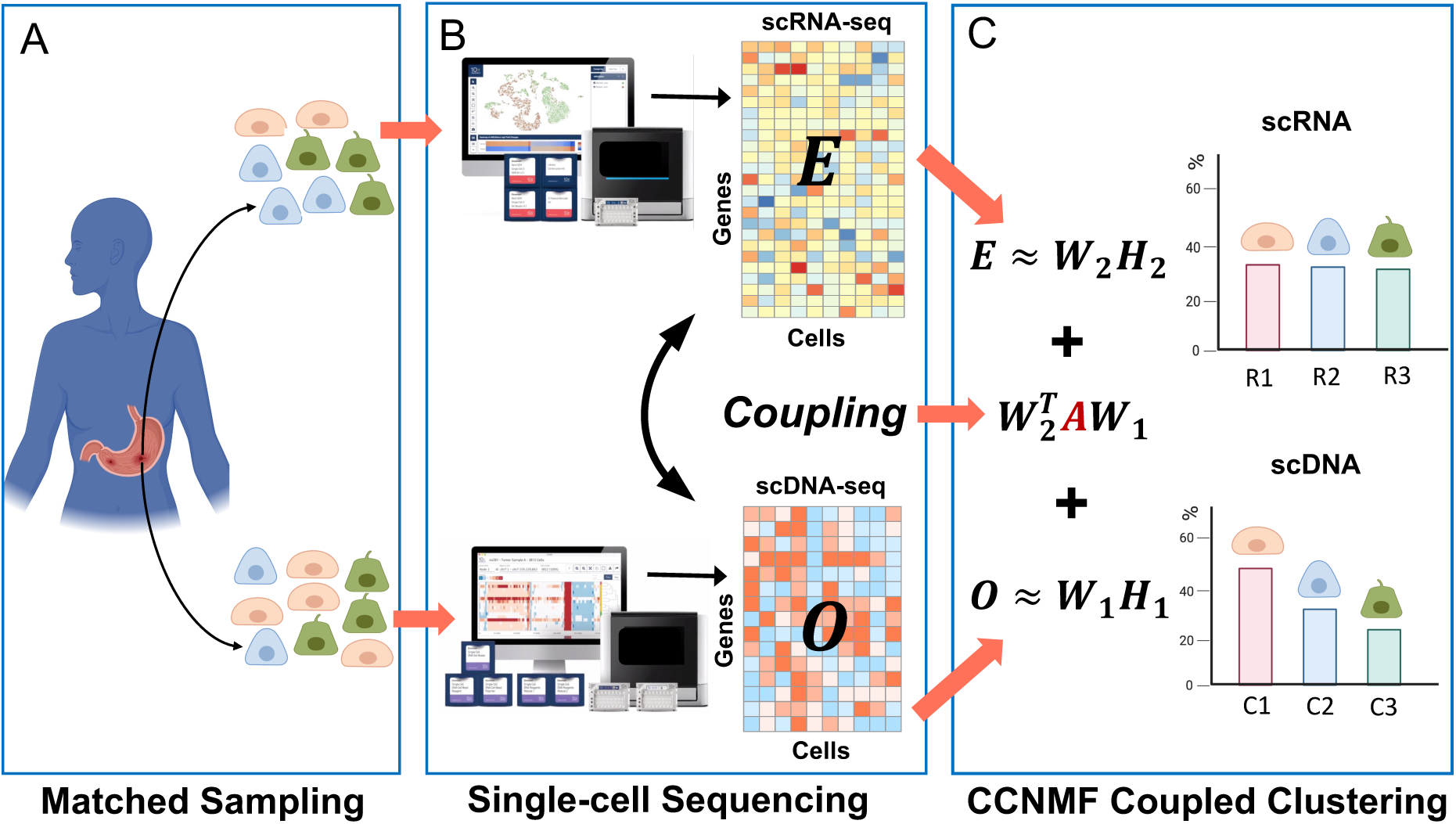
The workflow of Coupled-Clone Nonnegative Matrix Factorization (CCNMF). It took scDNA matrix *O ∈ R^p×n^*^1^ and scRNA matrix *E ∈ R^p×n^*^2^ as input for inferring shared clonal structure between two modalities from the same bio-specimen. We applied a coupling matrix *A ∈ R^p×p^* to link matrices *O ∈ R^p×n^*^1^ and *E ∈ R^p×n^*^2^, which represented the global concordance between DNA copy number and gene expression.

The outputs of CCNMF include: *W* matrices that represent the expression or copy number profile centroids of scRNA or scDNA clones; *H* matrices that represent the cell-wise membership weights toward clone for each of scRNA and scDNA cells. The toolkit is platform-independent and works with general R installation. It is made available as an open source software on Github with detailed readme, manual and examples.

### CCNMF recovers the underlying clonal structures in Simulation

We firstly evaluated the performance of CCNMF using simulated scDNA and scRNA datasets – *Sim*. The evaluation was based on Adjusted Rand Index (ARI), and the results were presented in **Supplementary tables S1-S3**. *Sim* included two different scenarios, each with three parameters. In each scenario, ARI was assessed by varying the parameter of interest over its range while keeping the others as default. The default parameters for copy number fraction, outlier percentage and dropout percentage were 50%, 0 and 0, respectively.

**Supplementary table S1** showed the simulation results for the linear and bifurcate clonal structure scenarios, with various copy number fractions ranging from 10% to 50%. It was defined as the percentage of genome undergoing copy number changes. As shown in the table, CCNMF was able to recover the exact underlying clonal structure with the highest accuracy (ARI=1) for all cases under both scenarios, except for one case with an ARI of 98%. With the decreasing of the copy number fraction, the clonal copy number difference becomes smaller, making it harder for CCNMF to resolve the clones accurately. The results demonstrated that such effect is only modest, as with only 10% of genome having copy number difference between the clones, CCNMF was still able to correctly uncover the underlying structure.

**Supplementary table S2** showed the simulation results with varying dropout percentages from 10% to 90%. Dropout percentage was defined as the percentage of cells with zero values for gene expression and copy number. The reason behind it could be the limited sensitivity of a technology or the gene was not present or expressed. Dropouts are very common in scRNA and scDNA data because of amplification bias and other random effects. In *Sim*, a dropout percentage at 10% means that 10% of all simulated gene expression or copy numbers were perturbed to be zeros. As shown in the table, CCNMF achieved high accuracy in recovering the underlying clonal structure for all cases under both scenarios. All resulted ARIs were *>* 98%, except for one case with ARI of 81%.

**Supplementary table S3** showed the simulation results for outlier percentage ranging from 10% to 90%. It was defined as the percent of cells with non-realistic copy numbers or expression values. In practice, these data points are typically deemed technical errors and are excluded from downstream analysis. In *Sim*, outlier percentage stands for the percent of all simulated scDNA and scRNA cells were perturbed to be outliers. It was obvious from the table that CCNMF was robust to the presence of outliers. When the outlier percentage was less than 60%, CCNMF achieved high accuracy with all ARIs greater than 92% for both scenarios. In summary, our comprehensive simulation study demonstrated that CCNMF achieved good performance for resolving the coherent underlying clonal structures in scDNA and scRNA data with practical noise considerations.

### CCNMF detected cell origins for ovarian cancer mixture cell lines ***OV*** data

We conducted CCNMF analysis on the *OV* benchmark dataset consisting of a mixture of ovarian cancer cell lines. It was composed of two cell lines, OV-2295(R) and TOV-2295(R) from the same patient. OV-2295(R) was an ascites site cell line that is abnormal adjacent tissue but not cancerous, while TOV-2295(R) was a high-grade serous ovarian cancer cell line (*28*). The ground truth for individual cells were obtained from the original publication (*28*).

CCNMF successfully characterized two subclones in the *OV* mixture cells lines. The scDNA cells were classified into subclones C1 and C2, which comprised of 394 cells from TOV-2295(R) and 371 cells from OV-2295(R), respectively. Correspondingly, the scRNA cells were categorized into subclones R1 and R2, consisting of 4918 cells from TOV-2295(R) and 1717 cells from OV-2295(R), respectively. Notably, C1 and R1 were from the same cell line TOV-2295(R), while C2 and R2 were from cell line OV-2295(R), indicating corresponding relationship with scDNA subclones with scRNA subclones. The tSNE plots of Fig. 2A and 2B demonstrated the clear separation of the two mixture cells lines in scDNA and scRNA, respectively.

**Fig. 2.**
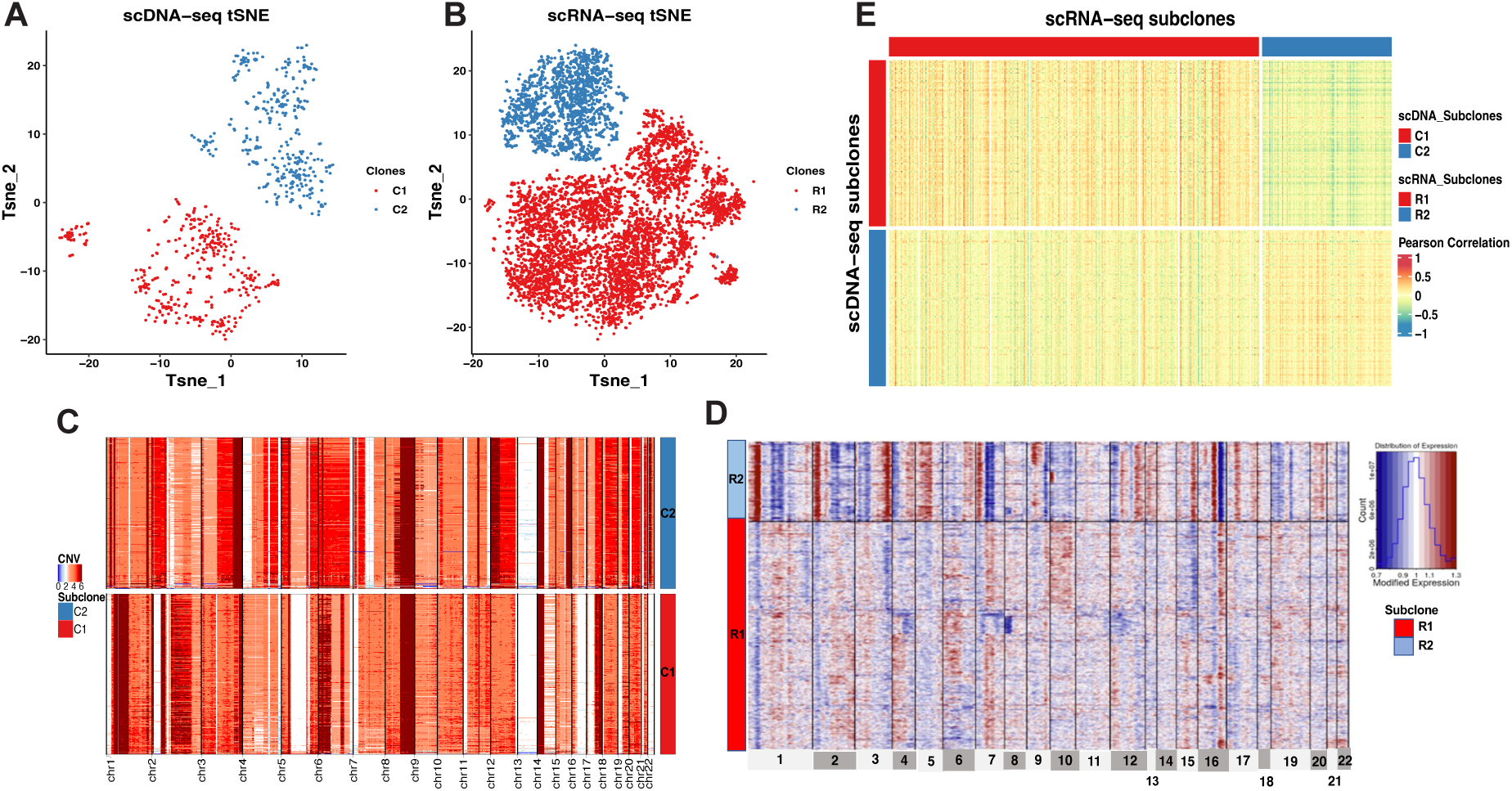
The coherent clonal structure between scDNA and scRNA of the mixture ovarian cancer cell lines. *OV*. (A) tSNE plot of scDNA clones. (B) tSNE plot of scRNA clones. (C) Heatmap shows CNVs changes across scDNA’s clones. (D) Heatmap shows inferred CNVs changes estimated by gene expression level across scRNA’s clones. (E) Heatmap of Pearson’s Correlation between single cells in coherent clones from scDNA and scRNA. Cells composing the same clone were coded in the same color. Each row represents a single cell and each column represents a genomic region for (C) and (D). The color in each dot of the heatmap represents the CNV status for (C) and (D). In panel (E), each row represents a single cell of scDNA, while each column represents a single cell of scRNA, the color in the heatmap represents the Pearson’s correlation of cells between scDNA and scRNA.

We compared the identified clones with the ground truth, resulting in an adjusted Rand index (ARI) of 1, indicating that CCNMF accurately recovered the underlying clonal structure of *OV*.

### CCNMF identified coherent clonal structure in gastric cancer cell line ***NCI − N87* data**

To investigate the performance of CCNMF on a cancer cell line, we analyzed the *NCI − N87* gastric caner cell line to determine whether the clone structure can be detected by CCNMF. The large-scale scDNA data of *NCI − N87* was composed of 1105 single cells. A substantial proportion of replicating cells existed in the scDNA which will cause analytical difficulty for clone reconstruction (*12*). It is because the most copy number changes in replicating cells were driven by ongoing DNA replication, which overwhelm the true clonal copy number variants. To address this challenge, we filtered out replicating cells before joint clustering analysis. We identified the replicating cells by calculating the variance of copy number for each cell because the higher replication activity gives rise to a higher variance of observed intra-cell segments. We utilized the Expectation-Maximization algorithm to fit a two-component mixture normal distribution on the obtained copy number variance across all cells (**Supplementary Fig. S4**). The cells were identified as replicating cells if they were assigned to a normal distribution with a larger mean, and were excluded from further analysis. Using this efficient filtering procedure, we successfully identified and removed the group of replicating cells with highly fluctuating copy numbers and retained 724 cells (**Supplementary Fig. S5**).

We utilized CCNMF to analyze single-cell copy number and gene expression matrices with selected and shared genes/features. We successfully identified and characterized three subclones (C1, C2 and C3) in the *NCI − N* 87 cell line for matched scDNA and scRNA, as shown in the tSNE plots (see Fig. 3A and 3B). Out of the initial 1005 cells of scDNA, 281 cells were identified as replicating cells and were discarded from downstream clonal reconstruction.

**Fig. 3.**
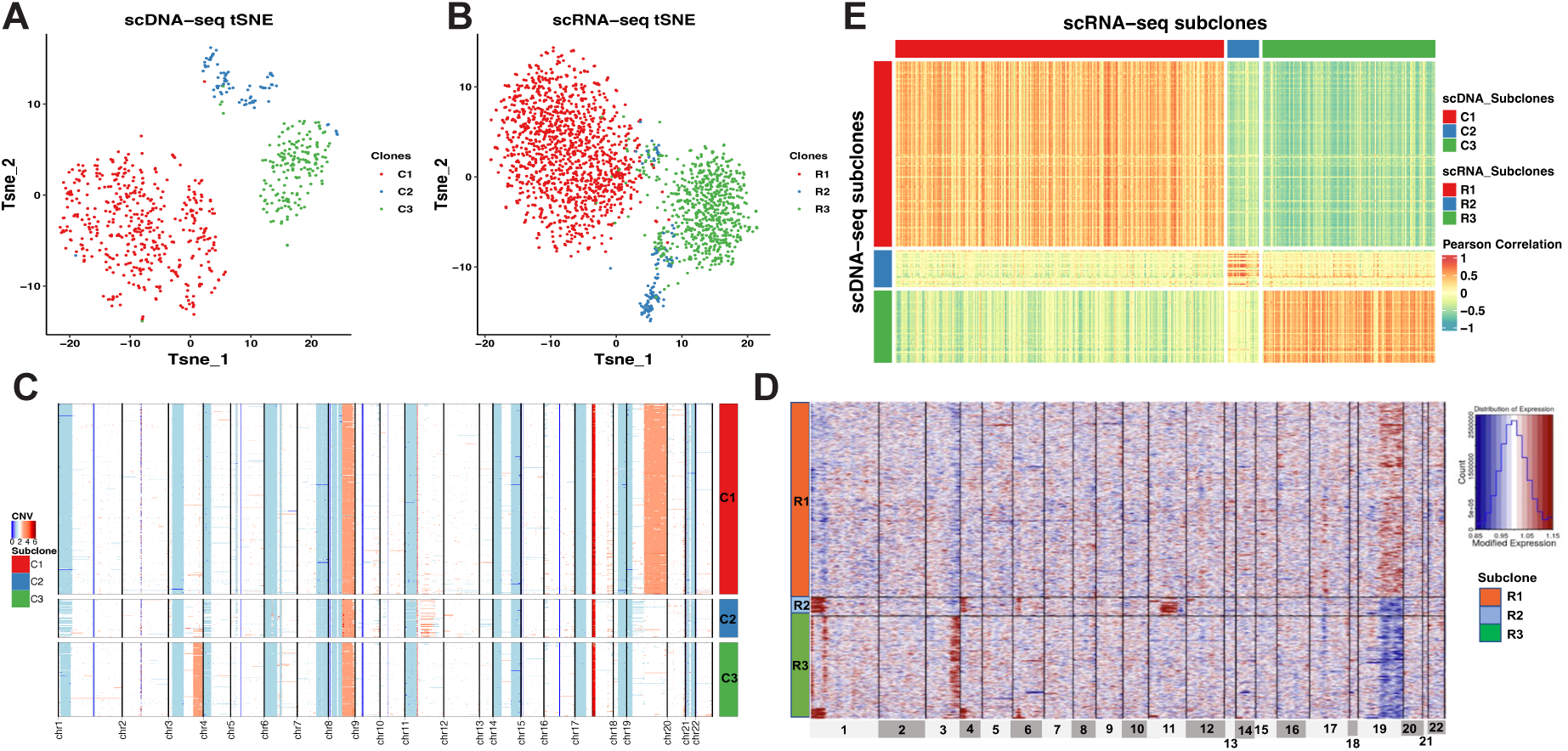
**The coherent clonal structure between scDNA and scRNA of the gastric cancer cell line** *NCI-N87*. (A) tSNE plot of scDNA clones. (B) tSNE plot of scRNA clones. (C) Heatmap shows CNVs changes across scDNA’s clones. (D) Heatmap shows inferred CNVs changes estimated by gene expression level across scRNA’s clones. (E) Heatmap of Pearson’s Correlation between single cells in coherent clones from scDNA and scRNA.

The remaining 724 cells were classified into three subclones, with 456, 91 and 177 cells in C1, C2 and C3, respectively. Notably, all the cells in C1 exhibited a consistent large CNVs amplification on the chromosome 19, while all cells in C3 shared a focal amplification event on the chromosome 3q arm. However, C3 was involved in a smaller proportion cells without amplifications on chromosomes 3 and 19, and instead, shared lesser amplification events on the chromosome 11 (see Fig. 3C). As an independent validation, our analysis successfully resolved the two major subclones and one minor subclones characterized in another study (*12*), with C1 and C3 corresponding to the two major subclones and C2 corresponding the minor one.

To increase comparability between scRNA and scDNA, we also filtered out the replicating cells in scRNA by calculating cell cycle scores using the “CellCycleScoring” function in Seurat. Ultimately, we selected 2168 scRNA cells in G0/G1 phase out of a total of 3246 cells and coupled them with 724 scDNA cells in G0/G1 phase as well. We identified three subclones (R1, R2 and R3) from 2168 scRNA cells, with 1337, 128, and 703 cells in each, respectively. To estimate the clonal large-scale CNVs, we applied the inferCNV package on gene expression with clone identities. We then visualized the inferred CNVs changes among the clones of scRNA by heatmap (Fig. 3D). Notably, the three scRNA subclones (R1, R2 and R3) corresponded to the scDNA clones (C1, C2 and C3). The inferred CNVs in R1 were consistent with C1, which shared a amplification event on chromosome 20. The focal amplification event on chromosome 3q arm was shared by R3 and C3. The cells of R2 shared lesser amplification events on chromosome 11, similar to C2. It is worth noting that the amplification-like events on chromosome 1 and 4 observed in R3 represented the relatively higher gene expressions in those cells than others in R1 and R2, which were potentially corresponding cells in C2 with diploids. We calculated Pearson’s correlation between pair-wise cells from scDNA and scRNA. It depicted a clear block pattern where cells in matched clones between two modalities had high correlations than cells in different clones, due to the clone-wise gene dosage effect associated with CNVs (Fig. 3E).

### CCNMF characterized cohort clone structure for primary gastric cancer *P5931*

Besides analyzing tumor cell lines, we performed CCNMF on real-world application for primary gastric adenocarcinoma. The scRNA and scDNA datasets were derived from a primary gastric cancer patient *P5931* (*29*). Primary tumors usually had higher degree of intracellular heterogeneity and more complex tumor microenvironment than cancer cell lines. Notably, we identified the tumor cells in G0/G1 phase for *P5931* scDNA after filtering out the replicating and normal cells (**Supplementary Fig. S6**). Moreover, we also detected the epithelial cells in G0/G1 phase in scRNA (*29*). Then we applied the CCNMF to find the underlying clone structure between tumor G0/G1 cells in scDNA and epithelial G0/G1 cells in scRNA for *P5931*.

Due to the heterogeneity in primary cancer specimen, there were no ground truth available of cell clone identities. Therefore, we interpreted the clonal structure based on the result of CCNMF. As shown in Fig. 4A and 4B, two clones overlaid with cell identities on tSNE plots for scDNA and scRNA. The two clones were clearly separated in scDNA through several cells in C1 were closer to C2 group. The clonal structure of scDNA of this primary gastric cancer sample was dominated by these significant copy number alternations affecting chromosomes 7 and 21 (Fig. 4C). The somatic events shared by cells of C1 were chromosome 8 and 21 amplifications. Clearly, the chromosome 7 amplification and chromosome 21 deletion were the major defining somatic events that separated the C2 from C1.

We visualized the underlying large-scale CNVs pattern of scRNA clones with cell identities from CCNMF (Fig. 4D). Here we applied the inferCNV package to infer the large-scale CNVs on the clones of scRNA. The inferred CNVs were estimated based on the relative gene expression levels among different clusters. In Fig. 4D, the two clones R1 and R2 of scRNA were corresponding with C1 and C2 of scDNA, respectively. Here, the inferred CNVs in R1 were consistent with C1, which shared chromosome 8 and 20 amplifications. Also chromosome 7 amplification and chromosome 21 deletion were shared by R2 and C2. The Pearson’s correlation was calculated between pair-wise cells between scDNA and scRNA. The correlation heatmap demonstrated that the cells in matched clones between two modalities had high correlations due to the clone-wise gene dosage effect associated with CNVs (Fig. 4E).

**Fig. 4.**
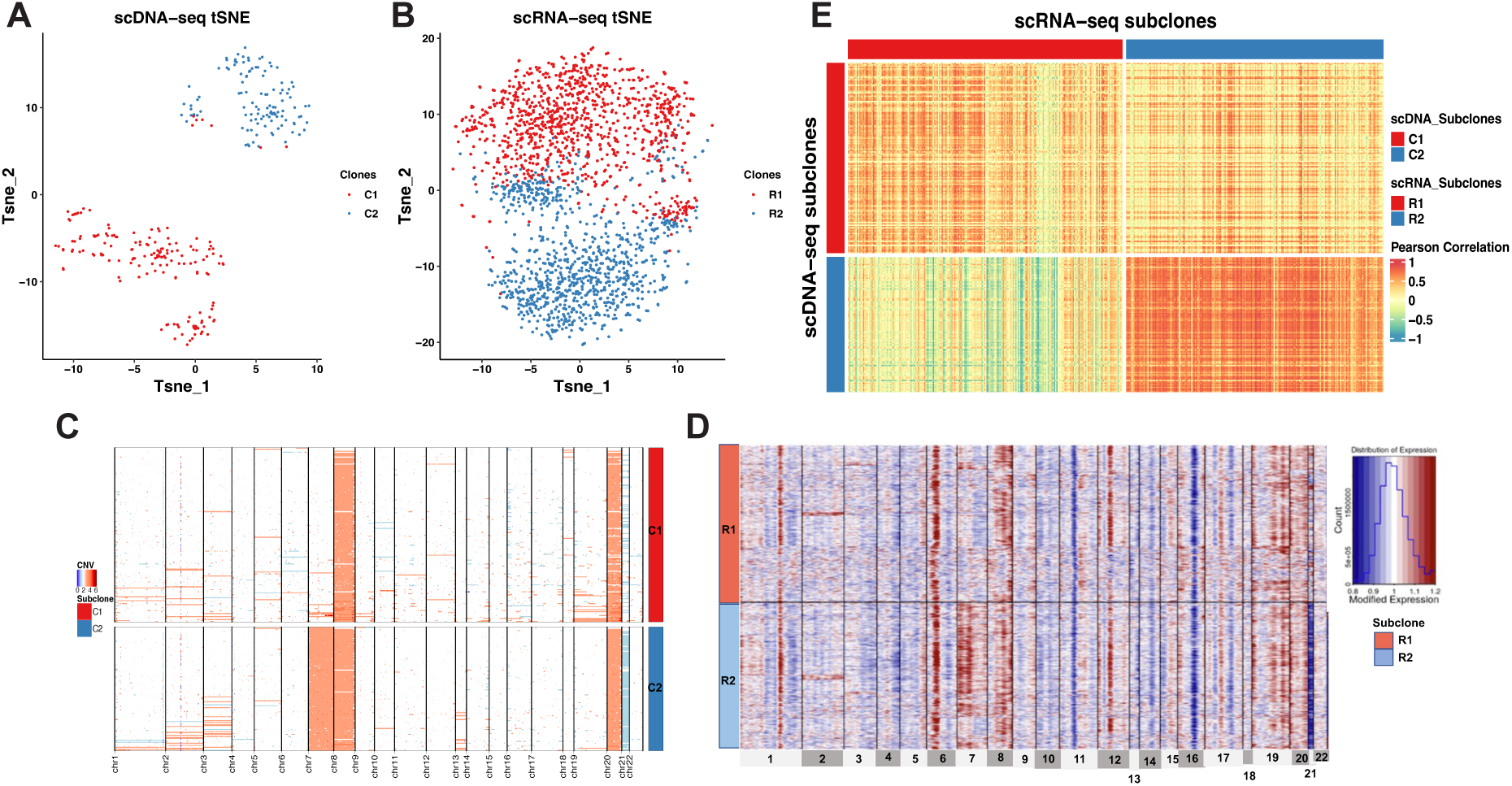
The coherent clonal structure between scDNA and scRNA of the primary gastric cancer patient. *P5931*. (A) tSNE plot of scDNA clones. (B) tSNE plot of scRNA clones. (C) Heatmap shows CNVs changes across scDNA’s clones. (D) Heatmap shows inferred CNVs changes estimated by gene expression level across scRNA’s clones. (E) Heatmap of Pearson’s Correlation between single cells in coherent clones from scDNA and scRNA.

## Discussion

Single-cell multi-omics technology enables the identification of cellular and genomic characteristics of cancer cells. The advancements in scDNA-seq technology allow for characterizing tumor subclone architectures by providing genomic DNA variation such as CNVs. Importantly, each subclone has distinct genomic alternations and cellular properties. Understanding the biological features and phenotypes of subclones is essential for precision cancer treatment. However, scDNA-seq data cannot be used to identify cellular types or phenotypes. Single-cell RNA-seq provides gene expression informations that elucidates the biology of individual cells, but it is limited in defining specific cancer subclones. Inferring CNVs from scRNA-seq remains as a potential area for obtaining biological insights of subclones (*20, 30, 31*). However, the inferred CNVs are based on changes in read depth or gene expression across the genome, and only provide limited reliability of copy number information of individual cells. Therefore, the matched scDNA-seq data is still needed as a ground truth of genome changes.

It is important to note that currently, there is a lack of technologies that are capable of simultaneously measuring copy number variants and gene expression of the same cell with high throughput. Although G&T-seq (*4*), DR-seq (*5*) and scTrio-seq (*6*) can measure genomic and transcriptome in up to a few cells per batch, there are low throughput manner. High throughput scRNA-seq can only measure transcriptome, and scDNA-seq is only for genome content, but not for both modalities in a single cell. To overcome these limitations, the integration of matched scDNA-seq and scRNA-seq of the same specimen is a promising approach.

To facilitate the understanding of tumor clonal structure and the associated biological characteristics like clonal-wise gene dosage effect, we proposed a joint-clustering approach CC-NMF for the integration of matched scRNA-and scDNA-seq data from the same specimen. CCNMF optimizes an objective function that simultaneously maximizes for intra-technology clonal compactness, inter-technology clonal coherence and expected dosage effect consistence. We demonstrated the utility of CCNMF by achieving high accuracy for resolving coherent clonal structure in simulation datasets with various noise levels. In real-world applications, CC-NMF has been validated by identifying the underlying clonal structure in mixtures of ovarian cancer cell lines, a gastric caner cell line and a primary gastric cancer patient. The consistent CNV change patterns between corresponding subclones between scDNA and scRNA were observed in results of CCNMF.

CCNMF uncover the coherent clonal architecture from the matched tumor single-cell genome and transcriptome data. The integration analysis showcased heterogeneous and clonal genetic nature of pathological tissues, which provides crucial dosage effect information for elucidating genetic cause and etiology of diseases. Furthermore, CCNMF can serve as a bioinformatics tool for performing single-cell level clonal dosage effect analysis for the community.

## Materials and Methods

### Coupled factorization of scDNA and scRNA data

We utilized the coupled-clone nonnegative matrix factorization framework to identify the underlying clonal structure of scDNA and scRNA data from the same biological specimen. The input can be any matched scDNA and scRNA metrics generated by various technologies. Input matrix *O ∈ R^p×n^*^1^ is the copy numbers of *p* genes and *n*_1_ cells from the scDNA; while the input matrix *E ∈ R^p×n^*^2^ is the gene expression of *p* genes and *n*_2_ cells from the scRNA (Fig. 1).

CCNMF was established on a powerful approach–Nonnegative Matrix Factorization (NMF), which uncovers the latent low-dimensional representation for a given matrix (*32, 33*). Briefly, NMF factorizes the given features by samples matrix into two non-negative matrices *W* and *H*, where *W* represents the latent structure of features (i.e. genes and CNVs), while *H* describes the weight of those features among samples (i.e. cells). Besides the NMF, the most important concept in CCNMF is to coupling the nonnegative factorization of matrices *O* and *E*. We additionally defined *A ∈ R^p×p^* to represent the linked sensitivity of gene expression to copy number. The matrix *A* serves as a bridge to enforce the link between changes in copy number and gene expression level for corresponding genes. *A* can be estimated priorly either by a linear regression model using public bulk RNA and DNA sequencing data (*22*) or by an identity matrix. The diagonal elements of the identity matrix represent the full informative links between copy number and gene expression on the same genes, but not for other genes. Hence, we simultaneously co-factorize the single-cell datasets *O* and *E* by minimizing the following objective function:

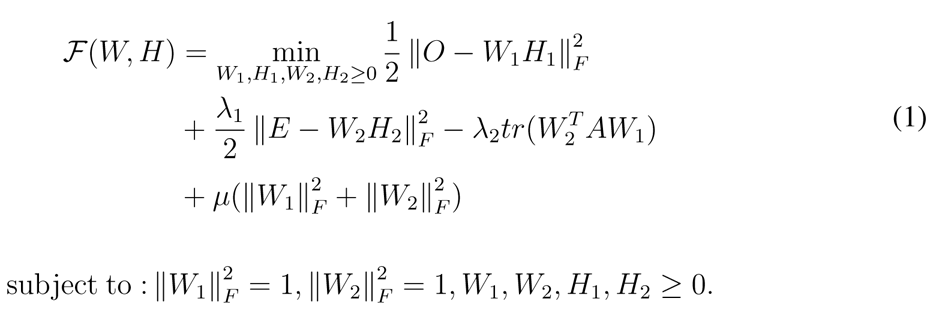

where we denoted *W*_1_ *∈ R^p×k^, W*_2_ *∈ R^p×k^* and *H*_1_ *∈ R^k×n^*^1^ *, H*_2_ *∈ R^k×n^*^2^ by shorthands *W* and *H*, *tr*() represents the trace of a matrix.

By minimizing the first two terms of the objective function in Equation (1), we ensured the respective NMF decompositions of *O* and *E* as *O* = *W*_1_*H*_1_ and *E* = *W*_2_*H*_2_. Notably,

*W_i_*(*i* = 1, 2) represents the mean matrix of clusters for the *n_i_*cells, while *H_i_*is the weight matrices that softly assign *n_i_* single cells to the underlying identity linked cell clusters. Upon convergence, the weight matrix *H_i_* provides the inferred cluster identities for all single cells by the maximum weight. Additionally, minimizing the cross term *−tr*(*W^T^ AW*_1_) ensured the coherence of the inferred clone structure between the scRNA and scDNA data.

### Optimization of the objective function

For optimization, we applied the alternating direction methods of multipliers (ADMM) (*34, 35*) to the objective function. Let *µ*_1_ and *µ*_2_ be the Lagrangian multipliers for *W*_1_ and *W*_2_, respectively, thus the transformed objective function was written as follows:

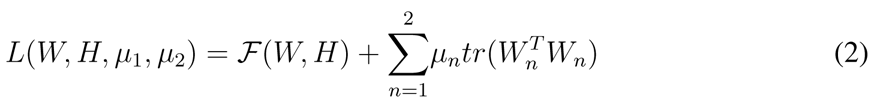

To solve the transformed objective function, we first obtained required gradients by setting its first order derivatives to zeros. Then, we used the gradient descent algorithm to iteratively update and optimize the objective function until convergence by following those steps (for proof see **Supplementary Methods**):

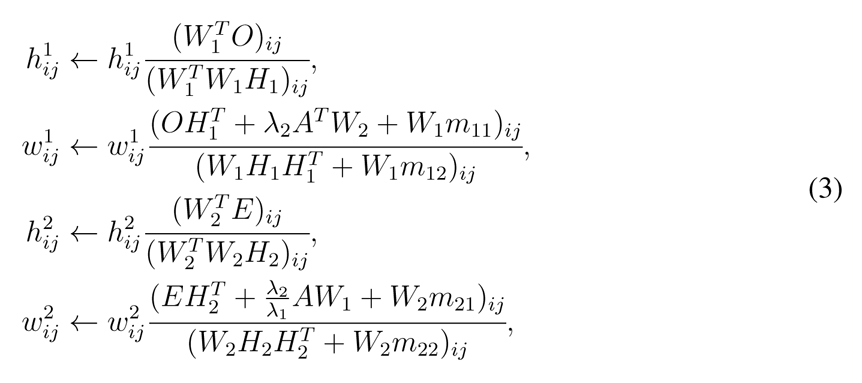

where

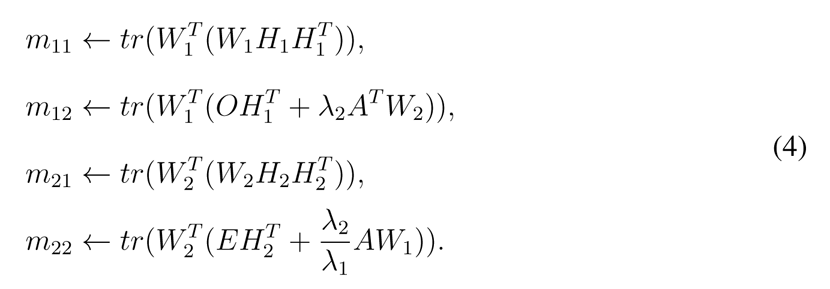

### Model inputs

The model had two hyper-parameters inputs as *λ*_1_ and *λ*_2_, which were used to initialize the iterative computation. Our experiences were that *λ*_1_ and *λ*_2_ are data-dependent. Nonetheless, they can be empirically determined by the input data. In practice, we used an automatic balancing strategy to determine the parameters, which ensured the initial values of the four terms of the objective function are within the same order.

The coupling matrix *A* is also expected as input, for which an identify matrix can be supplied. The non-zero diagonal elements represent the strengths of linked copy number and gene expression on the same gene, while the zero non-diagonal elements mean uninformative relationships between copy number and genes on different genes. To incorporate a real informative prior for *A*, we can estimate it from known associations between copy number and gene expression using bulk sequencing data of the same tissue source. It was well known that DNA copy number is highly positively correlated with the gene expression levels for the most (*>* 99%) of expressed human genes (*36*). We thus estimated a diagonal coupling matrix (*A ∈ R^p×p^*) where each diagonal element was calculated by the ratio of gene-wise mean expression to mean copy number using the bulk RNA and DNA data downloaded from The Cancer Genome Atlas (TCGA) (https://www.cancer.gov/tcga). Moreover, this obtained empirical matrix *A* also was used in simulation to generate realistic-like matched scRNA and scDNA datasets.

### Model Selection

When the number of cluster *k* is unknown, CCNMF runs specified range of models with different *k*. The optimal number of *k* was selected based on the minimum coverage value of the optimization function across various *k* sets.

### Preprocessing matched scRNA & scDNA data

We preprocessed the scRNA data with several steps, including filtering outlier genes and cells, normalizing for sequencing depth and log transformation. We also performed chromosome-level smoothing procedure for each cell, which the expression of order genes along each chromosome were smoothed the average of 101 gene window.

It is noted that scDNA features were located on the genome segmental bins, while scRNA features were presented across genes. To ensure the consistency of between scDNA and scRNA features, we associated multiple neighbor genome bins from scDNA with its corresponding gene from scRNA using the following preprocessing steps. First, we aligned both scRNA and scDNA into the same human genome assembly (GRCh37 or GRCh38). Then, we identified the one-to-multiple mapping between each gene and bins using the IRanges R package (*37*). Finally, the bin-level copy number variants of scDNA matrix was converted into the mapped gene by taking the average of CNVs on corresponding bins.

To perform feature selection on scDNA-seq copy number matrix, we developed a novel statistical approach due to the limited availability of methods in this task. After converting segmental bin-level scDNA data into gene-level CNVs, we selected the most highly variable genes across tumor cells while excluding replicating cells. Specifically, we modeled the variances of CNVs for all genes using a mixture normal distribution, and iteratively selected genes located at the normal distribution with a higher mean until at least 20% genes were chosen. The resulting genes were considered potentially informative and used as input features for CCNMF. Finally, we extracted the properly formed both scRNA and scDNA matrices *E ∈ R^p×n^*^2^ and *O ∈ R^p×n^*^1^ for CCNMF analysis (see Fig. 1), where *p* genes were the remaining selected features from scDNA.

### Simulation procedures

We generated the matched scRNA and scDNA data from the same clonal structure by presetting the ground truth genetic copy number (**GCN**) changes (as illustrated in **Supplementary Fig. S1**). Notably, the ground truth GCN profile represented a specific clonal structure. First, we specifically set the first clone as normal cells with GCN vector *V*_1_ = [2*, · · ·,* 2] *∈ R^m^*, where *m* enumerates over all genome segmental bins. The second tumor clone’s associated GCN vector as *V*_2_ *∈ R^m^*, in which a fraction of *V*_2_ were randomly sampled from *{*0, 1, 3, 4*}* with various probability parameters. Similarly, *V*_3_ of the third clone were also simulated by the above procedure. Finally, we obtained a ground truth GCN profile *V* = [*V*_1_*, V*_2_*, V*_3_*, · · ·*] as the baseline genomic change profile for the underlying clonal structure.

To simulate the scDNA data, we estimated their specific parameters and noise from bulk CNV datasets downloaded from cBioPortal (*38, 39*). Specifically, (i) we estimated the probability transition matrix *P* (*C|G* = *k*) for observed integer copy number when given genetic changes using bulk CNV data, where *C* is the observed CNV, *G* is the genetic CNV, *k* (0, 1, 2, 3*, · · ·*) is the value of copy number (**Supplementary Fig. S2**); (ii) We simulated copy number for per gene and per cell *D_ij_ ∼ multinomial*(*P* (*C|G*) *∗ P* (*V*)) (**Supplementary Fig. S3**) when given the clonal GCN matrix *V* ; (iii) We then added outlier and dropout events (**Supplementary Methods**).

We extended the associated genetic copy number (GCN) profile *V* to genes*×*cells matrix in which GCN determines the clone-wise genes expression and copies. We next used the Splatter pipeline (*40*) to simulate library effects, dropout, outlier events in scRNA-seq data based on the above extended clonal gene-wise matrix (**Supplementary Methods**). Splatter parameters were estimated from the same tumor tissue with RNA-seq and bulk CNV data downloaded from cBioPortal.

### Simulated and real datasets

As the first benchmark, we simulated 46 scDNA-and scRNA-seq datasets. They were referred as the **Sim** data. The linear and bifurcated clone structure scenarios with 3 clones were simulated in this study. We varied common experimental parameters, such as the percentages of outliers, dropouts, and CN dosage effect. For each simulated dataset, we randomly generated cell-wise scDNA and scRNA data according to the specified scenario and parameters using the procedure as detailed previously (also see **Supplementary Fig. S1**). Each of the obtained dataset in *Sim*, has 1000 cells and 2000 genes/CNV bins, and the three composing clones have 200, 400 and 400 cells each. The first clone was designed as normal cells, and the second and third clones represented by deletion and amplification events, respectively. We set the percentages of differentially acquired deletions and amplifications to affect 10% to 50% chromosome regions. We deposited the *Sim* data into GitHub.

For the real data benchmark, we obtained a mixture of high grade serous carcinoma (HGSC) cell lines for matched scDNA and scRNA, referred to as the **OV** data. It was sequenced by DLP scDNA-seq and the 10X Genomics scRNA-seq technologies and was downloaded from European Genome-Phenome archive with accession EGAD00001004553 (*21*). The mixture cells were made up from ascites (OV2295R) and solid tumors (TOV2295R). The scRNA subset included 1717 cells from ascites (OV2295R) and 4918 cells from solid tumors (TOV2295R), while the scDNA subset had 371 cells from OV2295R and 394 cells from TOV2295R.

As another real data application, we also downloaded the matched scRNA and scDNA data for *NCI − N* 87 gastric cancer cell line from Gene Expression Omnibus (GSE142750) and National Institute of Health’s SRA (PRJNA498809) (*12*). We firstly processed the scDNA-seq data using Cellranger-DNA pipeline with reference genome GRCh38 according to described procedures in (*12*). The scDNA data includes 1005 single cells, each contains 154423 copy number bins across the whole genome. The scRNA-seq data has 3246 cells with 13513 genes per cell.

Finally, we also collected a primary gastric adenocarcinoma patient data, called *P5931* (*29*), which was sequenced using 10X Genomics technologies for both scRNA and scDNA. The scDNA consists of 796 cells with 154423 copy number bins across the whole genome. The scRNA of *P* 5931 has in total 11217 cells with 19129 genes per cell. The data was downloaded from dbGAP repositories with accession numbers phs001711 (*29*).

## Acknowledgments

XB gratefully acknowledges the financial support from China Scholarship Council (No.201904910816).

## Author contributions

XB, LW and LCX designed the study; XB, ZD, LW and LCX were involved in algorithm development; XB performed data analysis; LW and LCX supervised this study; XB, LW and LCX wrote the manuscript. All authors reviewed and approved the final manuscript.

## Data and materials availability

The R package of CCNMF is available at https://github.com/labxscut/CCNMF.

## Supplementary materials

Supplementary Methods

Supplementary Figs. S1 to S6

Supplementary Tables S1 to S3

