## supplement for "Joint Inference of Clonal Structure using Single-cell Genome and Transcriptome Sequencing Data"

### Supplementary Methods

#### Proof about optimization for objective function

To couple the nonnegative factorization of matrices $O$ and $E$, we additionally define a matrix $A \in R^{p\times p}$ to represent the linked sensitivity of expression to copy number. The matrix $A$ can be estimated priorly either by a linear regression model using public paired RNA and DNA bulk sequencing data, or by using the uninformative identity matrix. Hence, we simultaneously co-factorize the datasets $O$ and $E$ by minimizing the following objective function

$$\mathcal{F}(W,H) = \min_{W_1,H_1,W_2,H_2\geq 0} \frac{1}{2}\left\|O - W_1H_1\right\|_F^2 + \frac{\lambda_1}{2}\left\|E - W_2H_2\right\|_F^2 - \lambda_2 tr(W_2^T A W_1) \tag{1}$$
$$subject\ to: \|W_1\|_F^2 = 1, \|W_2\|_F^2 = 1, W_1, W_2, H_1, H_2 \geq 0.$$

where we denote $W_1 \in R^{p\times k}, W_2 \in R^{p\times k}$ and $H_1 \in R^{k\times n_1}, H_2 \in R^{k\times n_2}$ by shorthands $W$ and $H$, $tr()$ is the trace of matrix.

Next, we applied the alternating direction methods of multipliers (ADMM) to find the gradients of Equation (2). Let $\mu_1$ and $\mu_2$ be the matrices containing the Lagrangian multipliers for $W_1$ and $W_2$, thus we had the transformed objective function as follows:

$$L(W,H,\mu_1,\mu_2) = \mathcal{F}(W,H) + \sum_{n=1}^{2}\mu_n tr(W_n^T W_n) \tag{2}$$

Then we will prove the update equations correctly are the optimal solution of the objective function (Equation 1). We just show the details about variable $W_1$, and the proof of $H_1$, $H_2$ and $W_2$ following the same procedure. As in Equation 2, the Lagrangian multiplier $\mu_1$ is for the constraint $\|W_1\|_F^2 = 1$. We simplified the Lagrangian function:

$$\mathcal{L}(W_1) = \frac{1}{2}\left\|O - W_1H_1\right\|_F^2 - \lambda_2 tr(W_2^T A W_1) + \mu_1(tr(W_1^T W_1) - 1) \tag{3}$$

Its first order derivative is:

$$\frac{\partial L(W_1)}{\partial W_1} = (W_1H_1H_1^T + 2\mu W_1) - (OH_1^T + \lambda_2 A^T W_1) \tag{4}$$

The KKT condition for the constranit $W_1 \geq 0$ gives

$$((W_1H_1H_1^T + 2\mu W_1) - (OH_1^T + \lambda_2 A^T W_1))_{ij} w_{ij}^1 = 0 \tag{5}$$

Combining with the constraint condition $\|W_1\|_F^2 = 1$, which means $tr(W_1^T W_1) = 1$, then

$$2\mu_1 = tr(W_1^T(OH_1^T + \lambda_2 A^T W_1) - W_1H_1H_1^T) \tag{6}$$

Consider the Equation (4) = 0 and Equation (6), we conduct

$$W_1H_1H_1^T - (OH_1^T + \lambda_2 A^T W_1) + W_1 tr(W_1^T(OH_1^T + \lambda_2 A^T W_1 - W_1H_1H_1^T)) = 0 \tag{7}$$

Finally, we get the update rule for $W_1$

$$w_{ij}^1 \leftarrow w_{ij}^1 \frac{(OH_1^T + \lambda_2 A^T W_2 + W_1 m_{11})_{ij}}{(W_1H_1H_1^T + W_1 m_{12})_{ij}} \tag{8}$$

where

$$
\begin{aligned}
m_{11} &\leftarrow tr(W_1^T(W_1H_1H_1^T)),\\
m_{12} &\leftarrow tr(W_1^T(OH_1^T + \lambda_2 A^T W_2)).
\end{aligned} \tag{9}
$$

Finally, we used the obtained gradients with a descent algorithm to iteratively update and optimize the objective function until convergence by the following steps:

$$
\begin{aligned}
h_{ij}^1 &\leftarrow h_{ij}^1 \frac{(W_1^T O)_{ij}}{(W_1^T W_1 H_1)_{ij}},\\
w_{ij}^1 &\leftarrow w_{ij}^1 \frac{(OH_1^T + \lambda_2 A^T W_2 + W_1 m_{11})_{ij}}{(W_1 H_1 H_1^T + W_1 m_{12})_{ij}},\\
h_{ij}^2 &\leftarrow h_{ij}^2 \frac{(W_2^T E)_{ij}}{(W_2^T W_2 H_2)_{ij}},\\
w_{ij}^2 &\leftarrow w_{ij}^2 \frac{(EH_2^T + \frac{\lambda_2}{\lambda_1} A W_1 + W_2 m_{21})_{ij}}{(W_2 H_2 H_2^T + W_2 m_{22})_{ij}},
\end{aligned} \tag{10}
$$

where

$$
\begin{aligned}
m_{11} &\leftarrow tr(W_1^T(W_1H_1H_1^T)),\\
m_{12} &\leftarrow tr(W_1^T(OH_1^T + \lambda_2 A^T W_2)),\\
m_{21} &\leftarrow tr(W_2^T(W_2H_2H_2^T)),\\
m_{22} &\leftarrow tr(W_2^T(EH_2^T + \frac{\lambda_2}{\lambda_1} A W_1)).
\end{aligned} \tag{11}
$$

#### Framework of simulation procedures

We first evaluated CCNMF using simulated paired scDNA and RNA data following the procedure as illustrated in Figure S1. The simulation principle is to coherently generate scRNA and scDNA data from the same ground truth genetic copy number and clonality while also allowing adding sequencing platform specific noises. To simplify the simulation, we set the total number of clones to be $k = 3$ in all simulated scenarios. We always specified that the first clone (cluster) as normal cells with a genetic copy number profile vector $V_1 = [2, \cdots, 2] \in R^m$, where $m$ enumerates over all genome segmental bins.

We specified the second cluster to represent clonal deletions. We obtained its associated genetic copy number vector $V_2 \in R^m$ by replacing fractional components of $V_1$ with the absolute copy number values randomly sampled from $\{0, 1\}$ according to parameters. Similarly, we specified the third cluster to represent clonal amplifications and obtained $V_3 \in R^m$ by replacing fractional components of $V_1$ with copy number randomly sampled from $\{3, 4\}$. We also recorded the ground truth clonal genetic copy numbers as $G_i^{CN}$.

Next, we defined the observed copy number per gene and cell as $O_i^{CN}$, which is the experimentally observed scDNA copy number data. We recognized that various batch, sequencing and platform noises can affect the genome segmentation results from experiments and cause $O_i^{CN}$ to deviate from $G_i^{CN}$. To realistically simulate $O_i^{CN}$, we used a Markovian model, which we estimated the transition probability matrix $P(O^{CN}|G^{CN})$ from the bulk copy number data of the TCGA project. To simplify the computation, the dimension of $P(O^{CN}|G^{CN})$ was set to $C_{max} + 1$, such that the copy number states can range from 0 to $C_{max}$ . In practice, we chose $C_{max} = 4$ as the maximum cut-off for copy number, which means any copy number larger than 4 (inclusive) were grouped into the state $C_{max}$.

Specifically, we estimated the transition matrix as follows: we downloaded the TCGA genetic copy number difference $G_{diff}^{CN}$ data from cBioPortal [1, 2] with 171 triple-negative breast cancer basal samples on paired bulk RNA-seq and DNA-seq data. where $G_{diff}^{CN} = \{-2, -1, 0, 1, 2\}$ and 0 means diploid/normal. We transformed $G_{diff}^{CN}$ to the integral copy number by $G^{CN} = G_{diff}^{CN} + 2$. We also downloaded the TCGA tumor purities for all samples from Butte et al.[3] and denoted them by $Purity = \{p_1, \cdots, p_n\}$. To estimate $P(O^{CN}|G^{CN})$: (1) We compensated for the associated purity and arrived at the raw copy number number $R_n^{CNV} = (2 \times C_{diff}^{NR})/p_n + 2$; (2) We grouped the $R^{CNV}$ according to their underlying genetic copy number $G^{CN}$ status (see Figure S2); (3) We fitted a Gaussian mixture model to each grouped $R^{CNV}$ (see Figure S3); (4) We calculated the $P(O^{CN}|G^{CN})$ by non-parametric binning of the histogram from the fitted Gaussian mixture per $G^{CN}$ status.

Note that $P(O^{CN}) = P(O^{CN}|G^{CN}) * P(G^{CN})$ is the empirically estimated multinomial probability vector we will use to simulate observed copy number $O^{CN}$ given the underlying genetic $G^{CN}$. We therefore simulated per gene per cell scDNA data $D_{ij}$ by randomly sampling from these multinomial distributions, such that $D_{ij} \sim multinomial(P(O^{CN}|G^{CN}) * P(V_i))$. As the last step, we added technology, batch and platform specific outliers and dropouts to the simulated scDNA data following the same procedure as for simulating the scRNA data that we described immediately below.

We simulated the scRNA data based on their associated clonal copy number profiles using the Splatter pipeline [4]. Specifically: (1) We simulated the $i$-th clonal gene expression background with multiplying the copy number profile $V_i$ by the dosage effect [5], such that the gene-wise expression mean of the $i$-th cluster is $\lambda_i^{'} = \lambda_i * V_i$; (2) We proportionally adjusted the gene-wise means for each cell using every cell's library sizes ($L_j$) which can be fitted by a log normal function with the estimated parameters from real data (see details in [4]), where $\lambda_{ij}^{'} = L_j(\lambda_i^{'}/\sum(\lambda_i^{'}))$; (3) We generated reads for each gene and each cell where their counts followed a Poisson mixture with an outlier component, such that $X_{ij}^{'} = 1_{ij}^{O}(X_{ij}^{O}) + (1 - 1_{ij}^{O})X_{ij}$, $X_{ij} \sim Poission(\lambda_{ij}^{'})$, $1_{ij}^{O} \sim Bernoulli(\pi^{O})$, $\pi^{O}$ is the probability of outlier occurrence, and $X_{ij}^{O}$ is the outlier's expression; (4) We simulated cell-wise gene dropout events by randomly replacing fractions of the generated gene expression with zeros, such that $G_{ij} = 1_{ij}X_{ij}^{'}$ mimicking a dropout effect $1_{ij} \sim Bernoulli(1/(1 + \lambda_i^{'}))$.

### Supplementary Figures

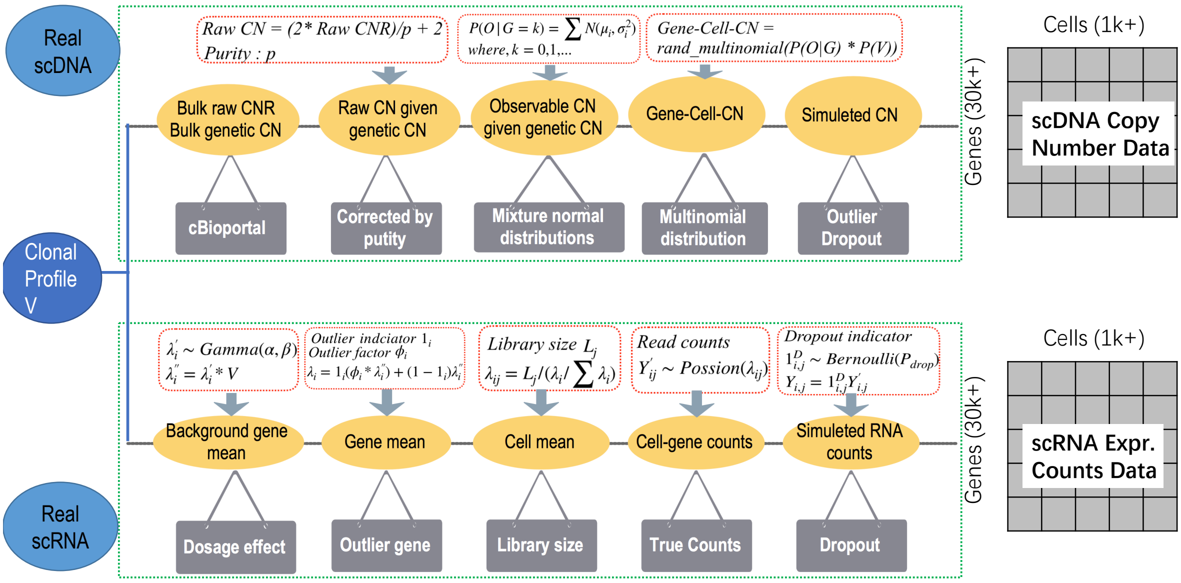


**Fig. S1.** The flowchart of simulation for paired single-cell DNA-seq and RNA-seq datasets. Firstly, we generate the associated clonal profile which has cluster/clone level's copy number change as **V**. For scRNA data, the parameters (as $\alpha, \beta, \phi_i, L_j, \lambda_{ij}$) can be estimated by Splatter pipeline from a real single-cell dataset. Then there are four procedures: (1) simulating the $i-th$ clone gene expression background by multiply the $i-th$ clone's CNV profile $V_i$ as dosage effect and generate the mean gene expression for $i-th$ cluster; (2) proportionally adjusting the gene means for each cell by the estimated library sizes; (3) generating read counts for each gene and each cell from a Poisson distribution by adding outlier; (4) simulating dropout events which each gene expression can be randomly replaced by zero according to *Bernoulli* distribution. As far as scDNA data, we first estimate the specific parameters of bulk copy number data from the same tissues with the above scRNA data. Thus, the bulk copy number ration difference and bulk genetic copy number can be downloaded from cBioportal (http://www.cbioportal.org). After that, (1) we estimate probability transition matrix after compensating the raw copy number variants with purity derived the observed; (2) grouping the RCNV according to their underlying different genetic copy number status (see Figures S2), and estimating the mixture normal distribution of each group row copy number as shown in Figures S3; (3) deriving the probability transition matrix by non-parametric binning of the histogram from mixture normal distribution given each genetic copy number status; (4) simulating the clustering CNV for each gene and each cell by randomly sampled from multinomial distribution; (5) simulating the outlier values and dropout events as the same as scRNA-seq simulator.

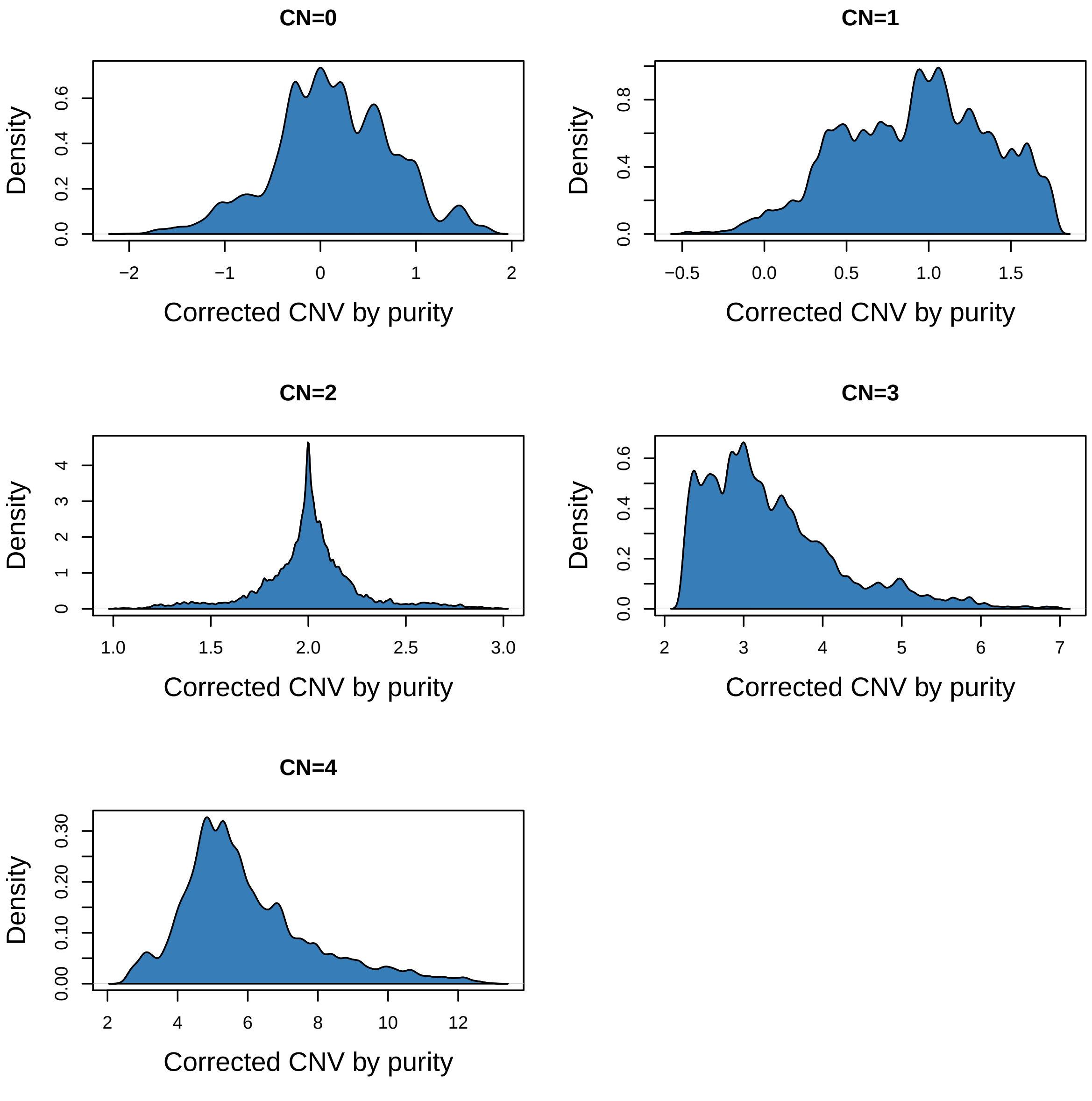


**Fig. S2.** The Fitted Observed gene-wise segmental copy number (CN) distributions by the underlying Genetic Copy Number Groups after purity correction.

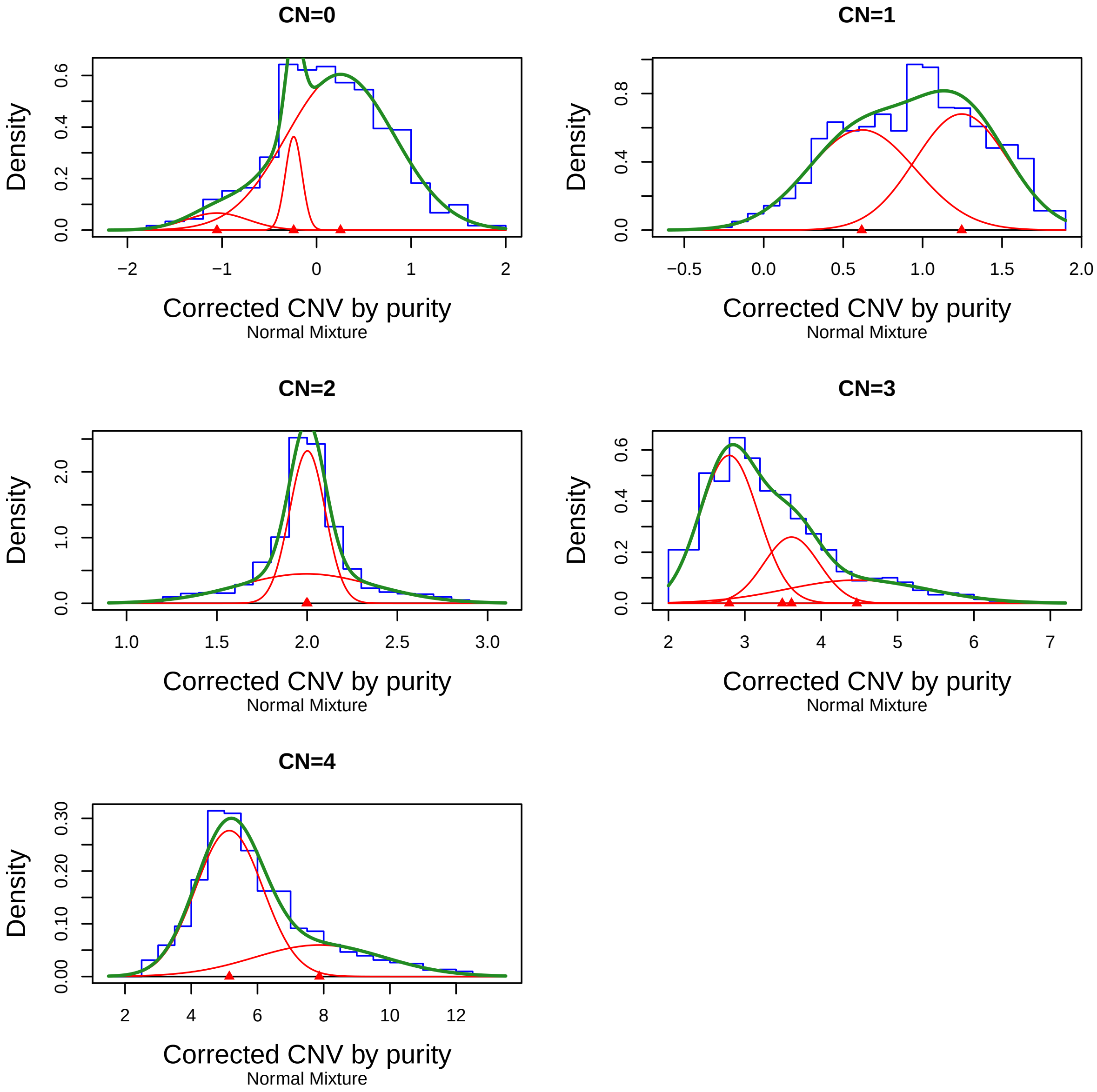


**Fig. S3.** The density of mixture normal distributions for Raw-CNV given genetic CNV (from 0 to 4). The blue line is the histogram density of Raw-CNV. The green line represents the density functions of fitted mixture normal model. The red lines are the normal distributions.

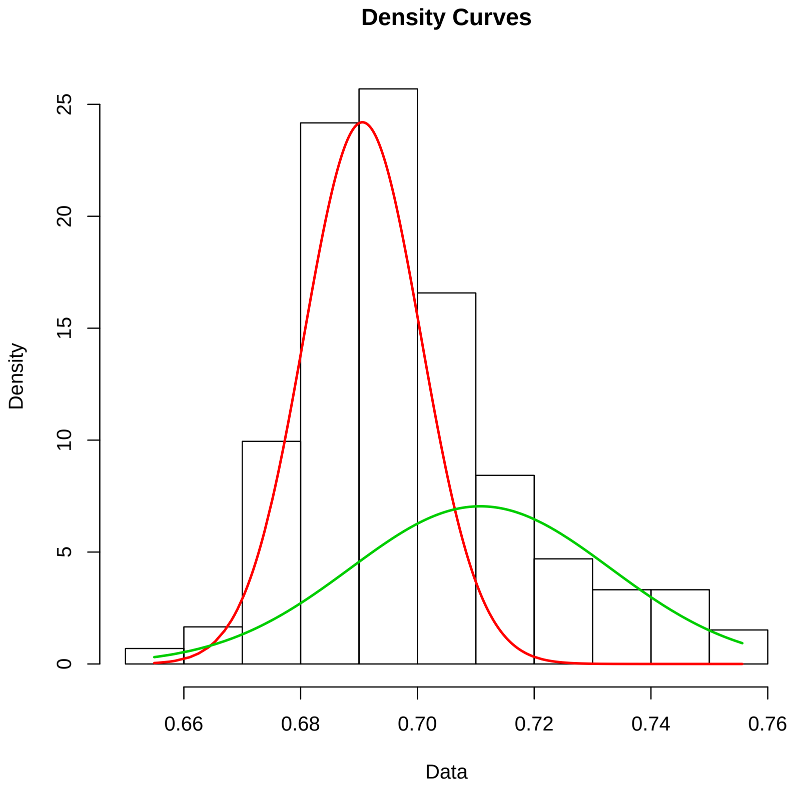


**Fig. S4.** The density of mixture normal distribution of cells' CNV variance. The cells belong to green part are replicating cells. y-axis is variance.

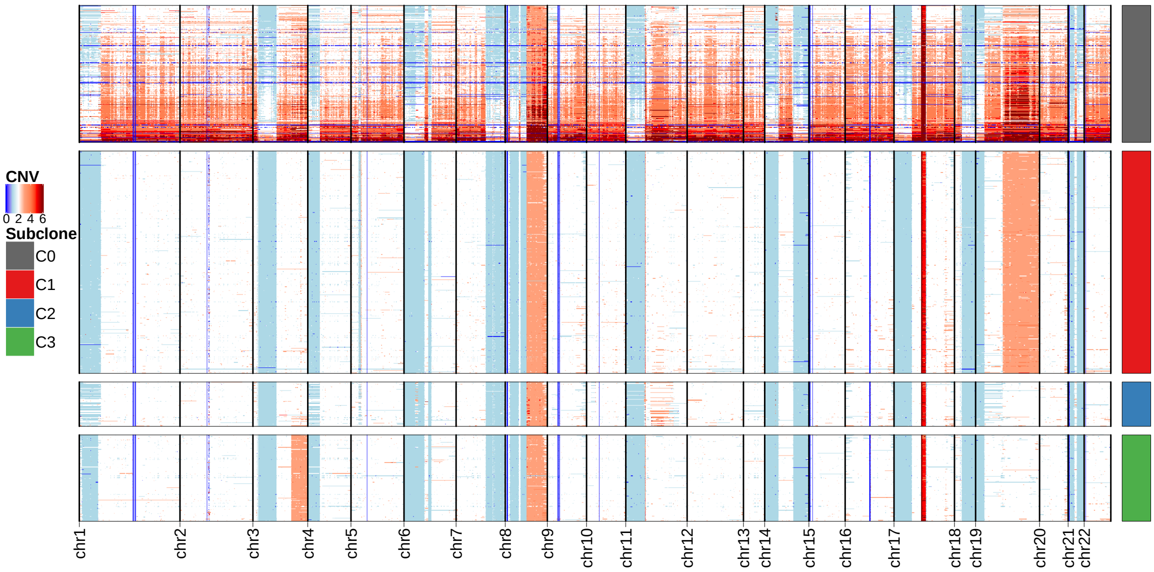


**Fig. S5.** Heatmap shows CNVs changes across replicating cells and scDNA's clones of *NCI-N87* cell line. Each row represents a single cell and each column represents a genomic region. The color in each dot of the heatmap represents the CNV status. C0 represents the identified replicating cells, C1 and C2 are the detected subclones of G0/G1 phase of scDNA.

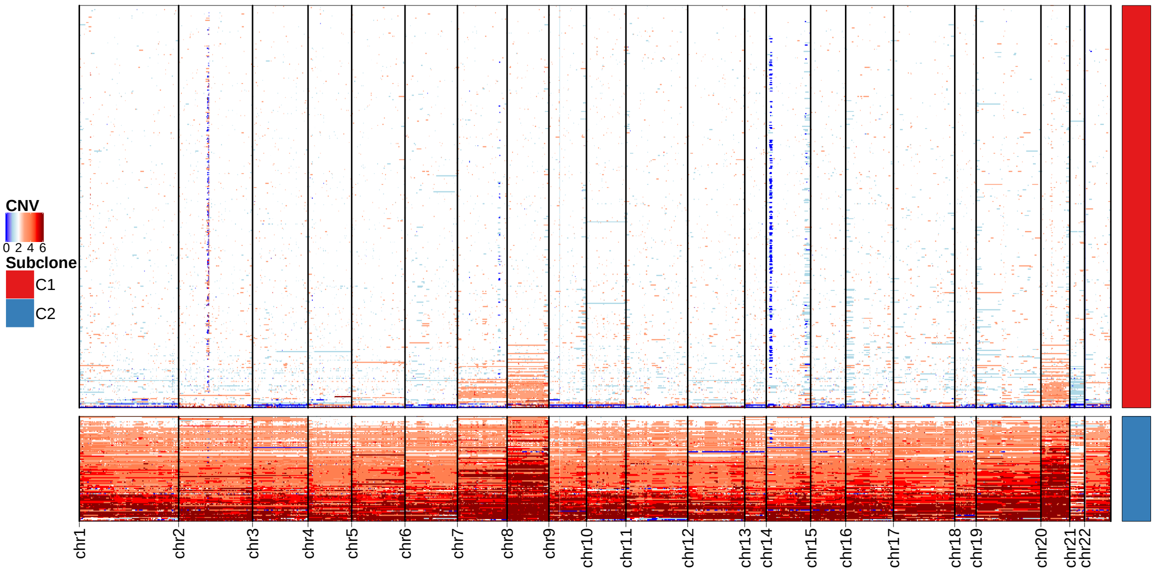


**Fig. S6.** Heatmap shows CNVs changes across normal cell and replicating cells *P5931* primary gastric cancer scDNA. Each row represents a single cell and each column represents a genomic region. The color in each dot of the heatmap represents the CNV status. C1 represents the detected normal cells, C2 represents the identified replicating cells.

### Supplementart Tables

Table S1. Performance (Adjusted Rand Index) of CCNMF by simulated copy number fractions

| Clonal Structure | Data Type | Simulated Copy Number Fractions | | | | |
|---|---|---|---|---|---|---|
| | | 10% | 20% | 30% | 40% | 50% |
| Linear | scRNA | 0.98 | 1 | 1 | 1 | 1 |
| | scDNA | 1 | 1 | 1 | 1 | 1 |
| Bifurcate | scRNA | 1 | 1 | 1 | 1 | 1 |
| | scDNA | 1 | 1 | 1 | 1 | 1 |

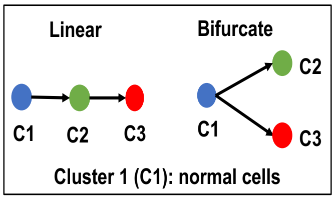


Table S2. Performance (Adjusted Rand Index) of CCNMF by simulated dropout percentages

| Clonal Structure | Data Type | Simulated Dropout Percentages | | | | | | | | |
|---|---|---|---|---|---|---|---|---|---|---|
| | | 10% | 20% | 30% | 40% | 50% | 60% | 70% | 80% | 90% |
| Linear | scRNA | 1 | 0.99 | 1 | 1 | 1 | 1 | 1 | 1 | 0.97 |
| | scDNA | 1 | 0.99 | 1 | 0.99 | 1 | 1 | 0.81 | 0.99 | 0.99 |
| Bifurcate | scRNA | 1 | 1 | 1 | 1 | 1 | 1 | 1 | 1 | 1 |
| | scDNA | 1 | 1 | 1 | 1 | 1 | 1 | 1 | 1 | 1 |

Table S3. Performance (Adjusted Rand Index) of CCNMF by simulated outlier percentages

| Clonal Structure | Data Type | Simulated Outlier Percentages | | | | | | | | |
|---|---|---|---|---|---|---|---|---|---|---|
| | | 10% | 20% | 30% | 40% | 50% | 60% | 70% | 80% | 90% |
| Linear | scRNA | 0.96 | 0.96 | 0.71 | 0.63 | 0.57 | 0.53 | 0.49 | 0.49 | 0.77 |
| | scDNA | 1 | 1 | 1 | 1 | 1 | 1 | 1 | 1 | 1 |
| Bifurcate | scRNA | 1 | 1 | 1 | 1 | 0.92 | 0.99 | 0.91 | 0.99 | 0.37 |
| | scDNA | 1 | 1 | 1 | 1 | 0.92 | 0.97 | 0.63 | 0.98 | 0.42 |

The performance of CCNMF on various configuration simulated datasets. There are main two structures including linear and bifurcate as shown in the left top box. For each structure, three configurations such as copy number fractions, outlier percentages and dropout percentages at different levels separately corresponds to Table S1, Table S2 and Table S3.
